# Adjacent Dorsolateral Prefrontal Cortex (DLPFC) Regions Participate in Distinct Large-Scale Networks Differentially Recruited for Social and Cognitive Control Functions

**DOI:** 10.64898/2025.12.22.695973

**Authors:** Lauren M. DiNicola, Randy L. Buckner

## Abstract

Human prefrontal cortex (PFC) is heterogenous. In monkeys, side-by-side PFC regions, including within dorsolateral PFC (DLPFC), show distinct long-range anatomical projection profiles, raising the possibility that adjacent regions might specialize as a consequence of the distributed networks in which they are embedded. Consistent with this possibility, recent findings in humans provide evidence for PFC regions domain-specialized for scene (spatial) processing and language processing that are adjacent to distinct domain-flexible regions responding to traditional cognitive control demands. Here we tested functional specialization of PFC regions linked to another domain-specialized network recruited by certain forms of social processing (theory-of-mind, ToM, tasks). Using within-individual precision neuroimaging approaches, side-by-side DLPFC regions embedded within parallel distributed networks were identified within the idiosyncratic anatomy of each individual (*N*=13). Functional responses between these distinct regions revealed a robust functional double dissociation: one DLPFC region was preferentially recruited by ToM tasks and an adjacent DLPFC region by working memory task demands. The region responding to social processing demands was small and its position varied slightly from one person to the next suggesting why it may have been underappreciated in past group-based analyses. These findings add to evidence that PFC features more functional specialization than commonly appreciated and further that the specialization of juxtaposed regions within PFC can be understood by examining the distributed networks within which the regions are embedded.

---

Prefrontal cortex (PFC) is associated with executive function and cognitive control (Milner & Petrides 1984, Goldman-Rakic 1987, Shallice & Burgess 1991, Miller 2000, Koechlin et al. 2003, Gilbert & Burgess 2008, Duncan 2010, Fuster 2015, Badre & Nee 2018, Mansouri et al. 2020 Friedman & Robbins, 2022). An unresolved question is how PFC is organized to support different functions. Does PFC comprise a hierarchy of interconnected regions that abstract and flexibly control broad domains of information processing or do PFC regions maintain domain specialization, playing distinct functional roles? As another alternative, does PFC organization possess elements of *both* domain-flexible and domain-specialized processing that might even operate in parallel? No single study or even line of studies will resolve these questions. Our strategy has been to chip away at them by estimating the organization of PFC using within-individual anatomical precision and then predicting and testing the functional response properties of PFC regions to constrain hypotheses about functional organization (DiNicola et al. 2020, Braga et al. 2020; DiNicola et al. 2023a, 2023b, Du et al. 2024). Here we extend these investigations by asking whether specific PFC regions participate preferentially in processing information within the social domain and whether specialization is predicted by the large-scale distributed network in which they are embedded. We focus on dorsolateral PFC (DLPFC).

In monkey neurophysiology studies, a subset of DLPFC neurons along the principal sulcus demonstrates delay activity during working memory tasks for combinations of object features (e.g., Rao et al. 1997, Rainer et al. 1998) and can flexibly shift what they represent dependent on the momentary task demands (Freedman et al. 2001). Similarly, human neuroimaging studies find that certain DLPFC regions respond when cognitive effort is increased across tasks that vary the types of information being processed (e.g., Duncan & Owen 2000, Fedorenko et al. 2013, DiNicola et al. 2023; see also Owen et al. 2005; Badre & Wagner 2007; Shashidhara et al. 2024). Motivated by these and related findings, a prominent model proposes that DLPFC functions as an integration zone, or hub, receiving convergent projections from domain-specialized brain systems and exerting broad top-down control on information processing (e.g., Miller et al. 2000, Miller & Cohen 2001). By this view, DLPFC should not possess domain-specialized processing regions.

An alternative hypothesis about DLPFC functional organization focusses on details of anatomical projection patterns and posits that distinct DLPFC regions maintain specialization for different processing domains (e.g., Goldman-Rakic 1988, Wilson et al. 1993). The key anatomical observation motivating this second view arises from analysis of single- and double-labeled tracer injections: neighboring posterior cortical regions project to distinct subregions of DLPFC, suggesting the presence of parallel, anatomically adjacent networks. By this view, distinct DLPFC regions maintain domain specialization and become differentially engaged dependent upon the information-processing domain required by a given task.

Findings from within-individual human precision neuroimaging studies raise the possibility that both ideas are correct with each describing distinct PFC regions, including within DLPFC itself, tightly adjacent to one another and easy to blur and confuse (e.g., Fedorenko et al. 2012, Michalka et al. 2015, Noyce et al. 2017, DiNicola et al. 2023a, Du et al. 2024, Assem et al. 2025, Ladwig et al. 2025). For example, Fedorenko and colleagues (2012) found that ventrolateral PFC (VLPFC) regions responding to high levels of cognitive effort are distinct from (but spatially adjacent to) regions preferentially responding within the domain of language. While it is possible that specialized regions within PFC act as lower-level regions that themselves project to higher-order integration zones (Miller, 2000), this possibility does not account for the differential response patterns that have emerged. Fedorenko and colleagues found that language-preferential VLPFC regions can be unaffected by increasing difficulty in a variety of working memory tasks and further that the nearby regions associated with working memory decrease their response when meaning-based language processing demands are increased. This functional double dissociation is hard to reconcile with a hierarchical model of prefrontal organization. Rather, the results are consistent with the possibility that multiple, parallel networks populate the association cortices, including network regions within PFC.

Functional specialization within PFC is not limited to language. We recently found another robust dissociation between DLPFC regions responding during remembering, specifically tracking scene (spatial) processing, and regions responding to domain-flexible cognitive effort (DiNicola et al. 2023a; see also Ladwig et al. 2025). Consistent with Goldman-Rakic’s parallel network model of PFC function, the response properties of spatially adjacent DLPFC regions were predicted based on the distinct large-scale networks in which they were embedded. Across numerous participants, a region of DLPFC just above the superior frontal sulcus responded robustly when the task demanded scene (spatial) construction but did not modulate to general working memory demands. This specific domain-specialized region of DLPFC is embedded within a distributed network involving parahippocampal cortex and hippocampus, suggesting its functional properties arise as an extension of the hippocampal formation’s role in memory and navigation. By contrast, a distinct adjacent region of DLPFC ventral to the superior frontal sulcus, participating in a parallel distributed network known as the “multiple-demand”, “frontoparietal control”, or “executive control” network (Duncan & Owen, 2000; Dosenbach et al., 2008; Vincent et al., 2008; Fedorenko et al., 2013), responded to increasing working memory load across both verbal and non-verbal domains. That is, the functional response properties of the side-by-side regions within DLPFC could be predicted based on the parallel networks in which they were embedded.

The present study takes this work further by exploring the social domain. Substantial work has identified brain regions that respond preferentially to social information processing, with particular emphasis on tasks targeting ‘mentalizing’ or ‘theory-of-mind’ (ToM; e.g., Fletcher et al. 1995, Saxe & Kanwisher 2003; see also Frith & Frith 1999, Koster Hale & Saxe 2013, Schurz et al. 2021, Frith & Frith 2021). Regions differentially responding to ToM tasks prominently include the temporal parietal junction (TPJ), superior temporal sulcus (STS), posteromedial cortex (PMC), and medial PFC (MPFC; e.g., Koster Hale & Saxe 2013; see also Van Overwalle 2009, Schurz et al. 2021). While MPFC is thus commonly studied in relation to social cognition (see also Amodio & Frith 2006, Lieberman et al. 2019), regions within lateral PFC, including DLPFC, are often conspicuously absent from description of the large-scale networks that preferentially process social information. Models of social reasoning that reference DLPFC often highlight a role in cognitive control separate from (or with top-down influence over) more specialized regions preferential to processing within the social domain (e.g., Lieberman 2007, Raz & Saxe 2020; see also Spreng & Andrews-Hanna 2015). An open issue is thus whether there exists PFC regions specialized for processing within the social domain that are distinct from PFC regions supporting domain-flexible cognitive control, as might be expected from a parallel network model of PFC organization (Goldman-Rakic 1988) and the general anatomical motif found for other association networks that consistently includes regions within PFC (Buckner 2026).

Preliminary evidence suggests specific PFC regions support processing within the social domain. We previously demonstrated and replicated that a large-scale distributed association network, termed Default Network B (DN-B), was preferentially active during ToM tasks and distinct from an adjacent network responding to tasks that place high demands on scene (spatial) construction (DiNicola et al. 2020). Critically, the functional double dissociation included regions in DLPFC (see DiNicola et al. 2020 Tables 6 and 7). In a recent comprehensive exploration of large-scale distributed association networks that examined multiple functional domains, Du et al. (2024) further found that DN-B could be functionally distinguished from an adjacent network supporting domain-flexible cognitive control, labeled Frontoparietal Network A (FPN-A). While the focus of the study was not on a specific zone of association cortex, domain-specialized regions of DLPFC were robust and present including for the social domain (see Du et al. 2024 Figures 32 and 33, and Figure A16; see also Ladwig et al. 2025).

Here, we build on these prior findings to formally test for a functional double dissociation between DLPFC regions responding to social processing demands as distinct from those linked to cognitive control. The locations and functional response properties of the specialized regions within DLPFC were predicted by the large-scale networks in which they are embedded, consistent with the parallel network model of PFC function presciently proposed by Goldman-Rakic (1988).

## Methods

### Participants

Data from the “Implementation Stage” participants reported in Du et al. (2024) were reanalyzed here, with participant numbers preserved. Thirteen right-handed paid participants with no history of neurological or psychiatric illness were included in analyses (18-34 years-old, mean= 22.5 yr, SD = 4.0 yr; 9 women; 6 self-reported as non-White and 3 as Hispanic). All participants provided written informed consent through protocols approved by Harvard University’s Institutional Review Board.

### MRI Data Acquisition and Processing

Data were acquired using a 3T Prisma^fit^ MRI scanner and 32-channel phased-array head-neck coil (Siemens Healthineers, Erlangen, Germany) at the Harvard University Center for Brain Science as described in Du et al. (2024). Each participant performed multiple scanning sessions (8-11 sessions each) that acquired resting-state fixation and task-based data using blood oxygenation level-dependent (BOLD) contrast. An openly available pipeline (‘iProc’; see also Braga et al. 2019), which maximizes cross-session alignment within individuals and minimize spatial blur and interpolation, was used to process the data. iProc integrates tools from FreeSurfer (v6.0.0; Fischl 2012), FSL (v5.0.4; Jenkinson et al. 2012) and AFNI (Cox 1996; 2012). All data were resampled to the fsaverage6 cortical surface mesh (using trilinear interpolation) and smoothed with a 2-mm full-width-at-half-maximum Gaussian kernel. Analyses were performed on the cortical surface within the idiosyncratic anatomy of the individual participants.

### Task Paradigms

Participants completed a battery of tasks as described in Du et al. (2024). Here we focused on the working memory (N-Back) task and two theory-of-mind tasks (False Belief and Emotional/Physical Pain Stories paradigms).

#### Working Memory (N-Back) Task

Participants each completed 8 runs of an N-Back task targeting cognitive control (4 min 44 s each; 37 min 52 s total; e.g., Braver et al. 1997; Cohen et al. 1994; see also Owen et al. 2005). The N-Back task varied working memory load via low-load (0-Back) and high-load (2-Back) conditions across four stimulus categories (Letters, Words, Scenes and Faces). Stimuli included consonant letters, one-syllable words (in sets of 10 matched for length and frequency using the Corpus of Contemporary English, Davies 2010), and color images (faces from the HCP – Barch et al. 2013; scenes from Konkle et al. 2010, Josephs et al. 2020). N-Back task parameters were modeled after the HCP (Barch et al. 2013). Each run featured 8 total blocks (one 0-Back and one 2-Back per category), each lasting 25 s (2 s condition cue and nine 2.5 s trials - 2 s stimulus and 0.5 s intertrial interval). Every second trial block was followed by a fixation block (15 s). Each trial block featured two target stimuli (i.e., matching the initial cue in the 0-Back condition or a stimulus seen two prior in the 2-Back condition). For Face, Scene and Letter categories, targets were identity matches; Word targets rhymed. Each block also included two lures (i.e., repeated or rhyming stimuli distinct from the cue in the 0-Back condition or in the wrong position in the 2-Back condition). Condition, category, target and lure positions were counterbalanced. Participants were instructed to respond “Match” (right pointer) or “No Match (left pointer) to each trial (see Du et al. 2024). Our primary contrast examined the load effect (2-Back vs. 0-Back) across all categories, with secondary analysis testing category-specific contrasts.

#### Theory-of-Mind (ToM) Tasks

Participants completed 8 total runs of ToM tasks targeting representation of others’ mental states (5 min 18 s each; 42 min 24 s total), with stimuli generously provided by the Saxe Lab (see Saxe & Kanwisher 2003, Dodell-Feder et al. 2011, Bruneau et al. 2013, Jacoby et al. 2016). Each run featured 10 unique trials (5 targeting ToM and 5 control), lasting 15 s each (10 s vignette, 5 s probe for response) and separated by 15 s of fixation. 4 runs of the False Belief task featured 10 unique stories about people or objects, followed by true/false statements requiring representation of either a person’s perspective (False Belief condition) or an object’s depiction (e.g., photos or maps - False Photo condition; Saxe & Kanwisher 2003, Dodell-Feder et al. 2011). Both conditions could include ‘false’ representations (e.g., incorrect beliefs or outdated information). 4 runs of the Emotional/Physical Pain Stories task featured 10 unique stories in which a main person experienced either emotional (Emo Pain condition) or physical (Phys Pain condition) pain. After each story, participants rated the protagonist’s pain on a 4-point scale from “None” to “A lot.” Our primary contrast averaged ToM contrasts from each task (False Belief vs. False photo and Emo Pain vs. Phys Pain), with secondary analysis testing task-specific ToM contrasts.

### Data Inclusion and Quality Control

Exclusion criteria were as described in Du et al. (2024) including maximum absolute motion >1.8mm and/or slice-based SNR<130, with a few lower SNR runs retained if they passed visual inspection. For the N-Back task, runs featuring more than two missed trials were excluded. For the ToM tasks, no runs featured more than one missed trial. Two participants were fully excluded from ToM analyses in Du et al. 2024 (P5 and P11); these participants were similarly excluded here. P3 had one run of the Pain task and P9 had one run of the False Belief task excluded, both due to motion. P10 had one run of the N-Back task excluded due to missing multiple trials.

### Network Estimation

Resting-state fixation task data were used for functional connectivity analysis and network estimation (with estimates carried forward from Du et al. 2024). Multiple resting-state fixation runs were acquired at each session (7 min 2 s per run, up to 24 runs total; see Table 1 in Du et al. 2024), with 16 (P2) to 24 (P12) total runs included per individual. Fifteen networks were estimated using a multi-session hierarchical Bayesian Model (MS-HBM; Kong et al. 2019, Du et al. 2024).

### Hypothesis-Targeted DLPFC Regions

Our primary aim was to test for a functional dissociation between nearby DLPFC regions embedded in two distinct large-scale networks. For the two networks of interest, DN-B and FPN-A, regions were identified within the MS-HBM outputs: a DLPFC region of DN-B and a nearby DLPFC region in FPN-A extending toward the inferior frontal gyrus. While each network’s regions are broadly localized to the same general vicinity and show similar relative positions, there was considerable variability in the precise region locations and boundaries across participants, including differences between the hemispheres. Example DLPFC regions in the left hemisphere are shown in Figure 1 (and see Supplemental Materials for all individuals’ bilateral regions). Primary analyses included bilateral regions; *post hoc* analyses investigated each hemisphere separately. For one participant (P6) there was no DN-B region identified in the right hemisphere and they were excluded for the hemisphere-specific analyses, but otherwise included in all primary analyses.

**Figure 1.**
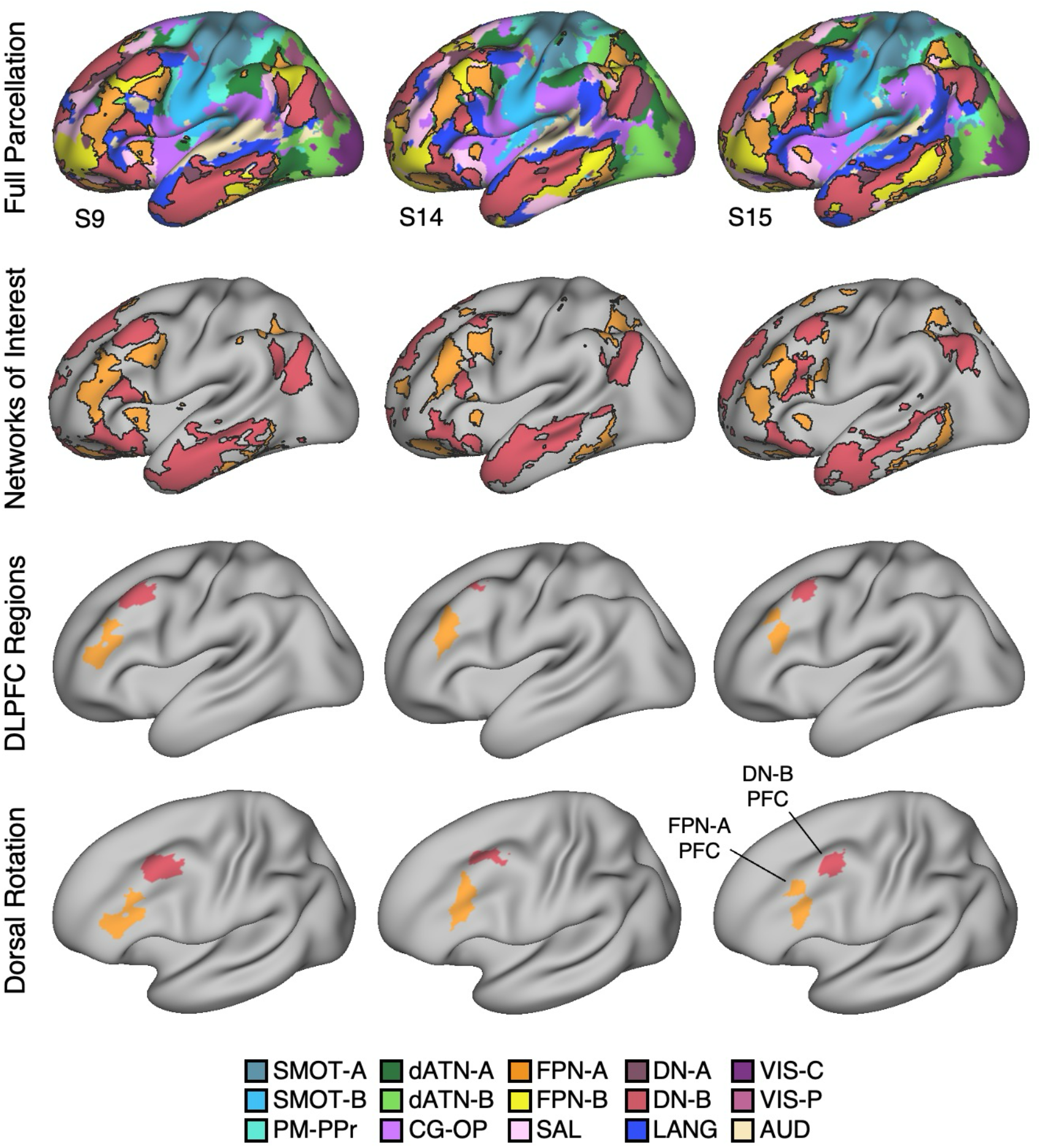
Dorsolateral prefrontal cortex (DLPFC) regions of two distinct networks identified within individuals using precision neuroimaging. Full network estimates and derived DLPFC regions are shown for three example individuals (one per column). Images display the left cerebral cortex, but regions were defined bilaterally. **(Top Row)** 15 networks were estimated in each participant using a multi-session hierarchical Bayesian model (MS-HBM). Colors correspond to network labels (shown in the bottom legend). Network estimates were taken directly from Du et al. (2024). **(Second Row)** The two primary networks of interest – DN-B (red) and FPN-A (orange) – were identified in all participants. **(Third Row)** Anatomically separate DLPFC regions of DN-B and FPN-A were isolated. **(Bottom Row)** A rotated surface images shows a complete view of the DLPFC. Network name abbreviations: Default Network A (DN-A) and B (DN-B), Frontoparietal Network A (FPN-A) and B (FPN-B), Somatomotor Network A (SMOT-A), B (SMOT-B), Premotor-Posterior Parietal Rostral (PM-PPr), Dorsal Attention Network A (dATN-A) and B (dATN-B), Visual Network Central (VIS-C) and Peripheral (VIS-P), Salience (SAL) network, Cingulo-Opercular (CG-OP) network (also referred to as the Action-Mode Network, AMN), Language (LANG) network, Auditory (AUD) network. See Supplemental Materials for bilateral network estimates and regions for all individuals.

### Testing the Hypothesized Double Dissociation

We hypothesized a functional double dissociation between DLPFC regions of DN-B and FPN-A, predicted to preferentially respond to the ToM task contrast and the N-Back Load Effect contrast, respectively. Repeated-measures ANOVA tested the 2 x 2 interaction, and planned post hoc comparisons tested each contrast (paired one-tailed *t*-tests, due to directional hypotheses).

Surface-projected task data were analyzed within individuals using run-level general linear models (GLM; first-level FEAT via FSL v5.0.4; see Du et al. 2024). Each GLM included a high-pass filter (100 s / 0.01 Hz cutoff) and a canonical (double gamma) hemodynamic response function and its temporal derivative. For the ToM tasks, each GLM produced vertex-level *z-*statistical maps for the contrast of interest, which were averaged across runs and across tasks for each participant (using fslmaths; Smith et al. 2004). For the N-Back task, each GLM output vertex-level *z*-statistical maps for each trial block, which were averaged by condition (0-Back or 2-Back) across runs. A difference between condition mean maps then produced a single cross-session contrast map per individual. For each individual, *z* values were then extracted from the mean ToM and N-Back contrast maps using the *a priori-*defined PFC regions of DN-B and FPN-A, producing two mean *z* values per contrast per individual.

### Examining the Robustness of the Double Dissociation

The ToM and N-Back task contrasts each included multiple task and stimulus conditions. The primary ToM contrast used two tasks that share targeting of others’ mental states (False Belief vs. False photo and Emo Pain vs. Phys Pain). To examine whether contrast-level differences might influence functional results, separate mean contrast maps for each ToM task were created and then re-tested for a functional double dissociation between DLPFC regions of FPN-A and DN-B. The primary N-Back contrast combined four stimulus categories (Faces, Scene, Letters, Words). In post-doc analyses, contrast maps for each N-Back stimulus condition were created and then explored to determine whether the effect generalized across stimulus categories.

## Results

### Adjacent DLPFC Regions are Specialized for Social Processing and Cognitive Control

DLPFC regions of DN-B and FPN-A showed a robust functional double dissociation. DLPFC regions of DN-B preferentially responded to the ToM task contrast and DLPFC regions of FPN-A to the N-Back Load Effect contrast. The hypothesized repeated-measures ANOVA showed a significant interaction between task contrast and DLPFC region (*F*(1,12) = 1117, *p* < 0.0001, see Figure 2), and paired *t*-tests supported significant differences between DLPFC regions for each task contrast, in the predicted directions (ToM: *t*(12) = 15.09, N-Back: *t*(12) = 13.59; both *p <* 0.0001). Thus, the full cross-over interaction was confirmed.

**Figure 2.**
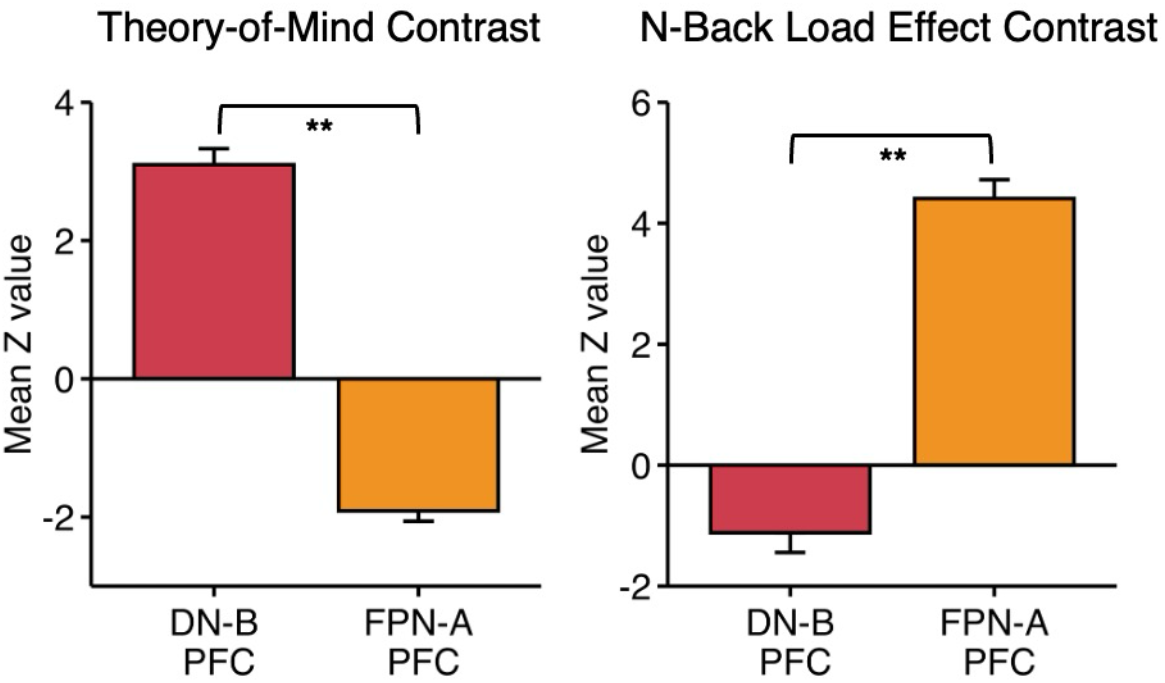
Adjacent regions of DLPFC demonstrate a robust functional double dissociation for Theory-of-Mind (ToM) and N-Back Load Effect contrasts. Panels show mean responses to ToM **(Left)** and N-Back Load Effect **(Right)** task contrasts. The ToM contrast preferentially recruits DLPFC regions of DN-B, while the N-Back Load Effect contrast shows a robust differential response in DLPFC regions of FPN-A. Plots show the means and standard errors of responses across individuals. The interaction effect was significant (*p* < 0.001), as were paired *t-*tests between regions for each contrast (** *p* < 0.001), indicating a full cross-over interaction.

DLPFC regions of DN-B were small bilaterally across participants, and for most the left hemisphere region featured more vertices than the right. *Post hoc* analyses restricted to each hemisphere revealed comparable results – interaction effects: Left *F*(1,12) = 441, *p* < 0.0001, Right *F*(1,11) = 818, both *p* < 0.0001; ToM: Left *t*(12) = 13.62; Right *t*(11) = 12.17, both *p* < 0.0001, N-Back: Left *t*(12) = 11.61; Right *t*(11) = 8.16, both *p* < 0.0001 (see Supplemental Materials). In analyses of right hemisphere data, P6 was excluded due to the absence of a detected DN-B region. All other participants were included in these analyses, including two additional participants for whom DN-B regions in the right hemisphere featured few vertices (P13 and P15).

### Functional Specialization Generalizes Across Tasks and Stimulus Conditions

We next examined whether the effects generalized across the included tasks and stimulus categories. Results aligned with the overall ToM patterns, with significantly greater recruitment of the DN-B regions of DLPFC for both False Belief and Pain task contrasts (False Belief: *t*(1,12) = 16.59; Pain: *t*(1,12) = 11.57; both *p* < 0.0001; see Figure 3). For the N-Back task, results from the four category-specific N-Back Load Effect contrasts all aligned with the overall N-Back Load Effect pattern, with significantly greater response by FPN-A regions than DN-B regions of DLPFC, regardless of category (Face: *t*(12) = 8.99; Scene: *t*(12) = 8.18; Letter: *t*(12) = 10.22; Word: *t*(12) = 6.21; all *p* < 0.0001; see Figure 4).

**Figure 3.**
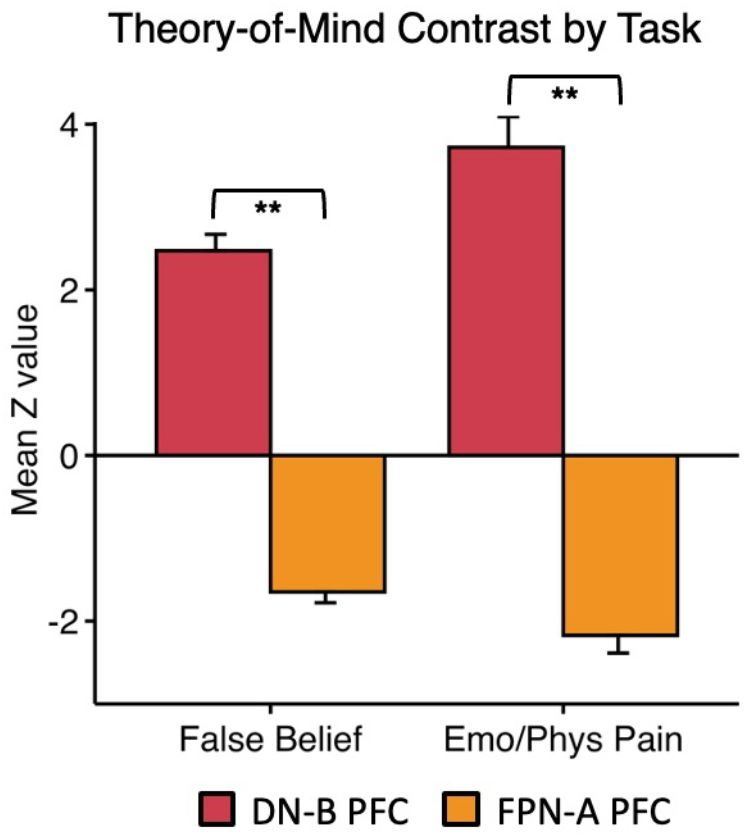
The DLPFC region linked to social inference responds across multiple Theory-of-Mind contrasts. Mean responses within DLPFC regions of DN-B and FPN-A to two separate Theory-of-Mind task contrasts – False Belief **(Left)** and Emotional / Physical (Emo/Phys) Pain **(Right)** – are shown. DLPFC regions of DN-B demonstrate a robust and preferential response in both task contrasts. Paired *t*-tests supported the observed functional dissociations (** *p* < 0.001).

**Figure 4.**
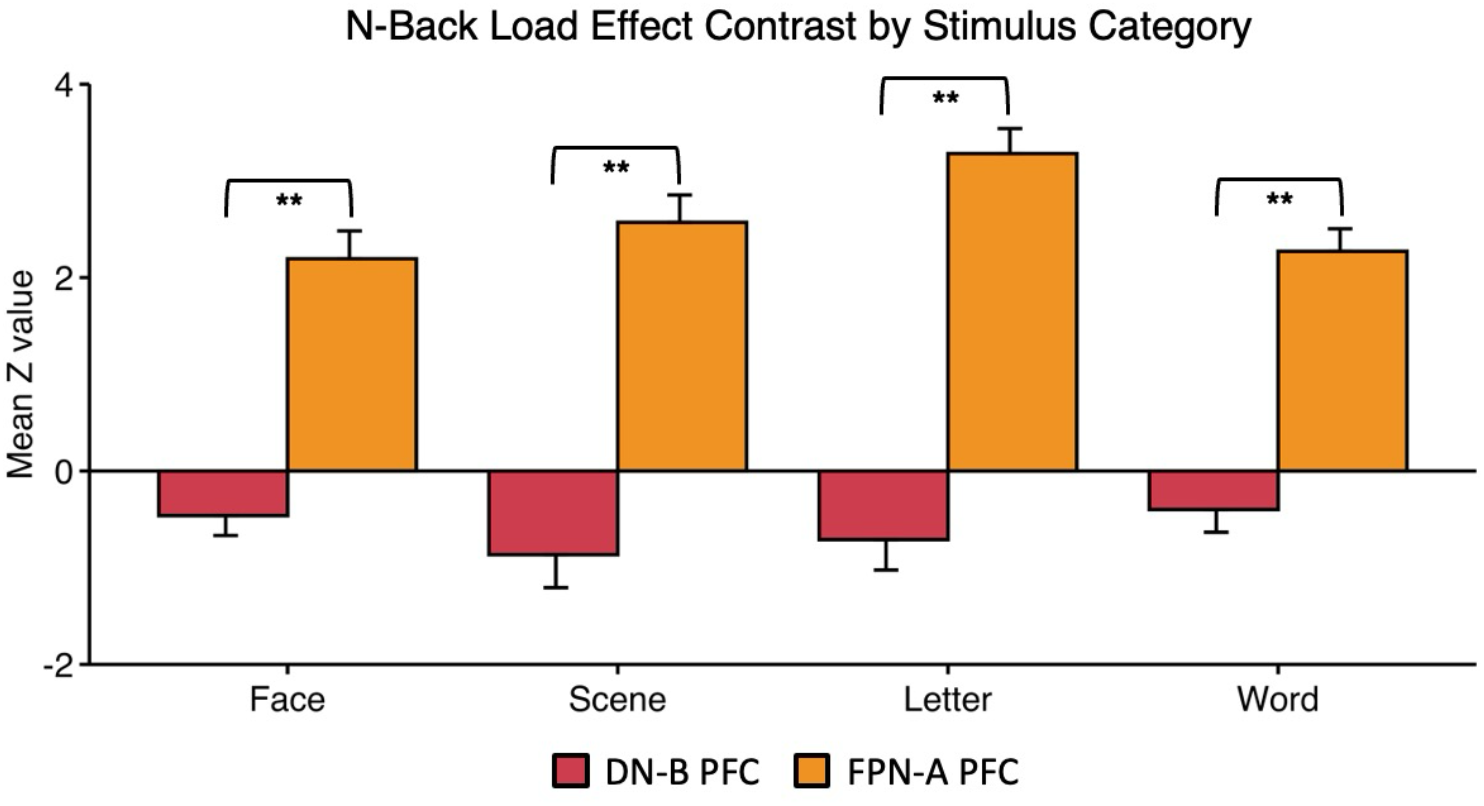
The DLPFC region linked to cognitive control responds flexibly across multiple stimulus domains. Mean responses within DLPFC regions of DN-B and FPN-A to the N-Back Load Effect contrasts across Face, Scene, Letter or Word stimulus conditions are plotted. As in prior work (DiNicola et al. 2023a), the regions linked to FPN-A show a strong response for all stimulus categories. Paired *t-*tests again supported the observed functional dissociations (** *p* < 0.001).

## Discussion

PFC features a mosaic of regions linked to distinct distributed networks, including within DLPFC. Here, we used precision neuroimaging to identify DLPFC regions embedded within two networks: a domain-specialized network that responds when making social inferences (called DN-B here), and a domain-flexible network involved in effortful control (FPN-A; e.g., Du et al. 2024; see also DiNicola et al. 2020, 2023). We found that DLPFC regions within DN-B responded to task contrasts involving the representation of others’ mental states. These domain-selective DLPFC regions did not increase response to traditional working memory demands. In contrast, adjacent, larger regions of FPN-A did show such an increase, consistent with a role in domain-flexible cognitive control (e.g., Cohen et al. 2000, Duncan 2010). These findings contribute to growing evidence that PFC is considerably more functionally specialized than commonly appreciated. Moreover, these specializations are predicted by a parallel network model of PFC organization: certain regions of PFC, including within DLPFC, maintain the functional properties of the large-scale distributed networks in which they are embedded (Goldman-Rakic 1988).

### DLPFC Contains a Domain-Selective Region Responsive to Processing of Social Information

Exploration of ‘social brain’ regions have often not included DLPFC (e.g., Mars et al. 2012, Koster-Hale & Saxe 2013, Paunov et al. 2019, Schurz et al. 2021). Even when analyses of social cognitive tasks report DLPFC response (e.g., in work on the canonical default network), the region is not emphasized as socially-relevant (e.g., Spreng & Andrews-Hanna 2015). Heuristically linking DLPFC to cognitive control has likely contributed to de-emphasis of DLPFC’s potential role in social processes. For example, proposals for why DLPFC might respond to social tasks include references to such control-relevant functions as increased effort and planning toward goals (e.g., during moral reasoning or in relation to others’ actions; see Forbes & Grafman 2010, Weissman et al. 2008) or ‘nonmentalizing aspects’ of social tasks (e.g., Lieberman 2007. These ideas may accurately reflect the roles of certain PFC regions that are embedded within domain-flexible control networks. The present findings suggest that DLPFC also possesses domain-specialized regions that participate preferentially in social processing functions relevant to ToM.

Across multiple studies, we have consistently observed that an anatomically specific distributed association network involved in social functions, termed DN-B, includes a small region within DLPFC (e.g., DiNicola et al. 2020, Du et al. 2024; see also Braga & Buckner 2017; Braga et al., 2019). The region’s location and extent vary from one person to the next, which likely contributes to difficulty in isolating its specialized functional properties, especially in analyses that use group averaging. When precision within-individual neuroimaging approaches are used, the extended network in which this DLPFC region is embedded is preferentially recruited by ToM task contrasts (DiNicola et al. 2020, Du et al. 2024, Edmonds et al. 2024). Here, we build upon those results, finding that DLPFC regions of DN-B (1) show preferential response to social over control task demands, (2) generalize their response across multiple forms of ToM task, and (3) are dissociable from adjacent DLPFC regions that increase response with cognitive load in traditional working memory tasks. While broad regions of PFC extending along inferior frontal gyrus do increase their response to effortful control, DLPFC regions of DN-B behave differently and preferentially respond to tasks that encourage processing within the social domain.

Future work will be required to better understand the anatomy and more detailed functional response properties of the domain-specialized regions of DLPFC. Edmonds and colleagues (2024) replicated topographic features (including DLPFC) and ToM response within DN-B and further described clear connectivity between this network and the amygdala (particularly basolateral and medial nuclei; 2024). Similar to our previously proposed hypothesis that a DLPFC region constitutes a component of a hippocampal-linked network involved in remembering and scene construction (e.g., DiNicola et al. 2023a; see also Angeli et al. 2025), perhaps a distinct DLPFC region is a component of a broader, domain-specialized, amygdala-cortical network that, in primates and especially so in humans, has evolved for higher-order processing within the social domain (see also Deen & Freiwald 2022).

### Growing Evidence Supports that Side-by-Side Prefrontal Regions Differentially Respond to Domain-Specific Processing and Domain-Flexible Cognitive Control

An ongoing debate about PFC purports either domain-flexible or domain-specialized regions. The present results build evidence for an alternative model: domain-specialized regions and domain-flexible regions *both* occupy LPFC, side-by-side (see also Fedorenko et al. 2012; Michalka et al. 2015; DiNicola et al. 2023a; Assem et al. 2025; Ladwig et al. 2025). As proposed by Fedorenko and colleagues (2012), regions responding in a domain-specialized manner (in their initial work, to language-relevant tasks) can be identified adjacent to regions responding domain-flexibly to tasks requiring effortful control. Crucial to that original and the present work, within-individual methods allow for identifying functionally dissociable regions, tightly juxtaposed and often spatially blurred in studies that rely on group averages or that otherwise blur across potential regional boundaries.

In a recent study, we replicated results from Fedorenko et al. (2012) that inferior frontal regions preferentially responding to sentence processing juxtapose broader regions recruited for domain-flexible control (DiNicola et al. 2023). Both sets of regions coupled to distributed, similarly dissociable association networks. We further found a DLPFC region specialized for mental scene construction, aligned with its associated, hippocampal-linked network (DiNicola et al. 2023a). That study thus showed multiple domain-specialized regions adjacent to a broader domain-flexible region in PFC. Here, we tested another piece of the PFC puzzle, showing that DLPFC regions of a third domain-specialized network - this one responding to socially-relevant tasks - can also be functionally dissociated from nearby regions participating in cognitive control functions. These ideas align with evidence from direct anatomical work in monkeys showing distinct prefrontal areas are interconnected in large-scale distributed circuits with different, juxtaposed areas of parietal cortex, forming parallel networks that support functional specialization (Goldman-Rakic 1988; see Buckner 2026 for discussion).

## Conclusion

Adjacent regions within DLPFC, identified through their participation in distinct distributed networks, respond preferentially to tasks involving theory of mind or cognitive effort. These results raise the possibility that an anatomically specific region of DLPFC may be part of an amygdala-linked network specialized for socially relevant processes. Most broadly, the present findings reveal that within PFC, multiple distinct and differentially specialized regions support either domain-specialized or domain-flexible control processes, in alignment with the broader networks in which they are embedded.

## Data Availability

Brain maps relevant to the present analyses are openly available (i.e., full network parcellation, SNR, task maps; https://balsa.wustl.edu/study/zK166). Code is also available (https://doi.org/10.7910/DVN/AVB4BW). Data from individual participants is available at the NIH repository (https://nda.nih.gov).

## Acknowledgements

We thank J. Du, P Angeli, N. Saadon-Grosman, W. Sun, S. Kaiser, J. Ladopoulou, A. Xue, B.T.T. Yeo, M Eldaief, T. O’Keefe, R. Mair, K. Ntoh, F. Davy-Falconi, and A. Billot for contributions to this project. We thank the Harvard Center for Brain Science neuroimaging core and FAS Division of Research computing, and R. Saxe, T. Konkle and E. Fedorenko for task stimuli. The multiband EPI sequence was generously provided by the CMRR at the University of Minnesota.

## Author Contributions

L.M.D. and R.L.B. conceived and designed the research. L.M.D. performed analyses. L.M.D. and R.L.B. interpreted results, prepared figures, and drafted, edited, revised and approved the manuscript.

